# Cognitive Flexibility Deficits in Rats with Dorsomedial Striatal 6-OHDA Lesions Tested Using a 3-Choice Serial Reaction Time Task with Reversal Learning

**DOI:** 10.1101/2020.04.08.022269

**Authors:** Zhuo Wang, Ilse Flores, Erin K. Donahue, Adam J. Lundquist, Yumei Guo, Michael W. Jakowec, Daniel P. Holschneider

## Abstract

Lesions of the dorsomedial striatum elicit deficits in cognitive flexibility that are an early feature of Parkinson’s disease (PD), and presumably reflect alterations in frontostriatal processing. The current study aimed to examine deficits in cognitive flexibility in rats with bilateral 6-hydroxydopamine (6-OHDA) lesions in the dorsomedial striatum. While deficits in cognitive flexibility have previously been examined in rodent PD models using the cross-maze, T-maze, and a food-digging task, the current study is the first to examine such deficits using a 3-choice serial reaction time task (3-CSRT) with reversal learning (3-CSRT-R). Although the rate of acquisition in 3-CSRT was slower in lesioned compared to control rats, lesioned animals were able to acquire a level of accuracy comparable to that of control animals following 16 days of training. In contrast, substantial and persistent deficits were apparent during the reversal learning phase. Our results demonstrate that deficits in cognitive flexibility can be robustly unmasked by reversal learning in the 3-CSRT-R paradigm, which can be a useful test for evaluating effects of dorsomedial striatal deafferentation.

## Introduction

Deficits in cognition, ranging from mild cognitive impairment (MCI) to dementia, are debilitating nonmotor symptoms in Parkinson’s disease (PD) [1]. Prior even to the appearance of motor symptoms, patients may manifest an impairment of executive function. Executive dysfunction in PD can elicit deficits in attentional control, cognitive inhibition, inhibitory control, working memory, and cognitive flexibility, all of which can impair a patient’s ability to plan, organize, initiate, and regulate goal directed behavior. This can lead even in early phases of the illness to difficulty in multitasking, initiating new tasks, and switching tasks. While the etiology remains to be fully understood, the frontostriatal circuits, dopaminergic and cholinergic systems have been implicated in executive dysfunction [2].

Early diagnosis and intervention at the stage of MCI are believed to be critical for treatment. Animal models and behavioral tests that allow investigation of PD-related cognitive deficits are key to mechanistic research and preclinical testing of new treatments. However, studies in this area have been relatively few compared to research on motor deficits. A challenge is that cognitive tests in animals often rely on motor functions and motor impairment in many animal models of PD can therefore be a confound. Recent preclinical PD research using an animal model has explored the dorsomedial aspect of the striatum, a brain region associated with behavioral flexibility and cognitive switching [3]. Cognitive tests using touch screens that presumably require only limited locomotor activities have also be utilized with PD animal models [4].

The 5-choice serial reaction time task (5-CSRT), modeled after clinical tests, has been broadly used to study operant learning, impulse control, and visual attention in rodents [5-7]. Tsutsui-Kimura et al. [8] reported a 3-choice variation (3-CSRT) to shorten the training time for rats to reach learning criteria. The current study applied 3-CSRT with reversal learning (3-CSRT-R) to test cognitive flexibility in rats with 6-hydroxydopamine (6-OHDA) lesion to the bilateral dorsomedial striatum. Whereas past research has generally used lever press in the 5-CSRT, it has been shown that nose poke is an easier response to learn in rats [9]. We therefore chose nose poke, which has been used in more recent 5-CSRT studies [6,7]. These choices were made to lower the task difficulty so that lesioned animals can rapidly learn the task to a similar level as controls before the initiation of reversal learning with a rule change. Thereby, cognitive flexibility can be investigated in separation from other cognitive deficits such as operant learning, attention, and impulse control.

## Methods

### Animals

Experiments were conducted under a protocol approved by the Institutional Animal Care and Use Committee of the University of Southern California, an institution approved by the Association for Assessment and Accreditation of Laboratory Animal Care, as well as by the Animal Use and Care Review Office of the US Department of the Army, and in compliance with the National Institutes of Health Guide for the Care and Use of Laboratory Animals. Male Wistar rats were purchased from Envigo (Placentia, CA, USA) at age 8 - 9 weeks. Animals were housed under standard vivaria conditions in pairs on a 12-hr light/12-hr dark cycle (dark cycle 6 p.m. – 6 a.m.).

### Animal model and stereotaxic surgical procedure

The 6-OHDA basal ganglia lesion model is a widely accepted model of dopaminergic deafferentation, and parallels many pathophysiologic features of the human disorder [10]. Animals were about 10 weeks old at the time of surgery. The procedure was as described before with changes in the injection sites [11]. To prevent any noradrenergic effects of the toxin, animals received desipramine (25 mg/kg in 2 ml/kg bodyweight in saline, i. p., Sigma-Aldrich Co., St. Louis, MO, USA) before the start of surgery. They were then placed under isoflurane anesthesia (1.5% in 30% oxygen and 70% nitrous oxide) in a stereotaxic apparatus (David KOPF Instruments, Tujunga, CA, USA) and received injection of 6-OHDA (Sigma-Aldrich Co., St. Louis, MO, USA) at four sites targeting the dorsomedial striatum bilaterally (AP: + 1.5, ML: ± 2.2, DV: - 5.2 mm, and AP: + 0.3, ML: ± 2.8, DV: - 5.0 mm, relative to the bregma), which is the primary striatal sector targeted by the medial prefrontal cortex (anterior cingulate, prelimbic area) [12]. Injection of 10 μg of 6-OHDA dissolved in 2 μL of 0.1% L-ascorbic acid/saline was made at each site through a 10 μL Hamilton 1701 microsyringe (Hamilton Company, Reno, NV, USA) fitted with a 30-gauge, blunted needle, at 0.4 μL/min controlled by a Micro4 microsyringe pump controller (World Precision Instruments, Sarasota, FL, USA). After injection, the needle was left in place for 5 min before being slowly retracted (1 mm/min). Naïve rats were used for controls. Carprofen (2 mg in 5 g tablet, p. o., Bio-Serv, Flemington, NJ, USA) was administered for one day preoperatively and for two days postoperatively for analgesia. Animals were left to recover for 2 weeks for lesion to mature prior to behavioral testing.

### Food restriction

Food restriction was started 2 weeks after surgery and maintained throughout the experiment. Animals were brought to 85% of their baseline bodyweight in one week and were allowed to gain 5 g in bodyweight per week thereafter, with ad libitum access to water. Animals were fed after behavioral training. Body weights were recorded Monday-Friday, with meal size individually adjusted on a daily basis.

### Tyrosine hydroxylase immunostaining

TH immunostaining data were collected from animals (*n* = 8) in a pilot study. Rats were humanely anesthetized and subjected to transcardial perfusion with 100 ml of ice-cold saline followed by 250 ml of ice-cold 4% PFA/PBS. Brains were removed, transferred to the same fixative for 24 hours, and then immersed in 20% sucrose for 48 hours. After sinking, brains were flash frozen, mounted and cut at 25 μm thickness on a cryostat microtome in the coronal orientation throughout the entire anterior-posterior extent of the brain. Selective sections representing levels of the brain spanning the site of 6-OHDA in both the striatum and midbrain were subjected to TH-immunostaining. Sections were rinsed with TBS at RT for 30 min, and quenched with 3% H_2_O_2_ for 10 min at RT. After rinsing in TBS, slides were washed in TBS+0.2% Triton-X 100 (TBST-0.2%) for 60 min at room temperature, then blocked with 4% NHS/TBST-0.2% and incubated overnight at 4°C in primary antibody solution (1:2500 anti-tyrosine hydroxylase, clone LNC1, Millipore, Billerica, MA) in 2% NHS/TBST-0.2%. Sections were visualized using secondary antibody solution (biotinylated anti-mouse IgG, Vectastain Elite ABC kit) 1:1000 in 2% NHS/TBST-0.05%, 60 min at RT. After rinsing with TBS, sections were placed in ABC Reagent for 60 min at RT. Staining was developed with 3,3’-diaminobenzidine (Vector Labs DAB Peroxidase Substrate Kit), until optimal contrast on sections was achieved. Sections were then mounted, dried overnight, and dehydrated before being coverslipped.

Images of tissues were made using an Olympus BX-50 microscope at low magnification, and the lesion size of the striatum was estimated bilaterally in the digitized, thresholded images of each rat by manual tracing using ImageJ 1.52k (Wayne Rasband, National Institutes of Health). Certain anatomical landmarks were used in the selection of striatal sections to demarcate rostral (bregma + 2.28 to + 1.28 mm, genu of corpus callosum with lateral appearance of the anterior commissures), mid (bregma + 1.28 to + 0.36 mm, medially located anterior commissure), and caudal levels (bregma + 0.24 to - 0.48 mm, posterior tail of the anterior commissure). Lesioned striatal areas were evaluated as a percentage of bilateral total striatal area.

### Spontaneous locomotor activity

Locomotor activity of the animals was recorded prior to 6-OHDA lesioning and two weeks thereafter. Recordings were performed in the vivarium during the dark cycle (6 p.m. – 6 a.m.). Animals were individually placed into clear plastic filtertop cages (46 cm length x 25 cm width x 22 cm height) with fresh direct bedding. During the recording period, animals were given a water gel cup for fluid intake but no food chow. Activity counts of each rats were recorded in the horizontal and vertical planes in time bins of 15 minutes by an infrared beam break system (Opto-M3, 160 Hz beam scan rate, 2.5 cm sensor spacing, 16 x16 sensor grid, Columbus Instruments, Columbus, OH) mounted around their cages. Data were collected from animals (*n* = 13) in a pilot study.

### Accelerating rotarod test

The effects of lesioning on coordination, balance and strength was evaluated on rotarod, a rotating cylinder treadmill with a diameter of 7.3 cm [11]. Rats were familiarized with the rotarod (Columbus Instruments, Columbus, OH, USA) at 2.3 m/min for 3 mins twice the day before testing. Rats were run using an acceleration paradigm (initial speed: 5 rpm = 1.15 m/min, acceleration rate: 6 rotations/min^2^ = 1.38 m/min^2^, 2 trials/day, 30-min intertrial interval for 2 days) until they fell onto a padded surface or reached the 5 min cutoff time (maximum speed: 35 rpm = 8.02 m/min). The outcome variable was the latency to fall averaged over four trials. Data were collected from animals (*n* = 8/group) in a pilot study.

### Sucrose preference test

We examined the effects of lesioning on sucrose preference to investigate anhedonia, which could emerge in toxin-induced models and therefore impact the motivation for reward in 3-CSRT. The protocol used was similar to those we have previously published [13]. To minimize neophobia, rats were exposed to the sucrose solution overnight for two days. The water bottle in each home-cage was replaced with two 50-ml bottles fitted with ball-point drinking spouts containing 2% sucrose. On the day of testing, rats were water-deprived for approximately 9 hours. At the onset of the dark phase, rats were individually housed overnight with access to 2 bottles, one containing 2% sucrose and the other water. Each filled bottle was weighed before and after the sucrose preference test, with fluid consumption measured by the difference. The location of the sucrose bottle (to the right or left side of the cage) was alternated to minimize side preferences. Sucrose preference was calculated as a percentage of total fluid intake, i.e. 100 x volume sucrose intake/ (volume of sucrose + volume water). Data were collected from animals (*n* = 8 lesioned, *n* = 6 controls) in a pilot study.

### 3-Choice serial reaction time task with reversal learning (3CSRT-R)

We modified the well-established 5-CSRT protocol and its 3-CSRT variation (Tsutsui-Kimura et al. 2009) (**Fig. 1A**). 1) Animals were trained through 3 difficulty levels with progressively shortened stimulus durations. While most 5-CSRT protocols train the animal at each difficulty level for a variable number of days until the animal reaches certain performance criteria, we chose to control the number of training days for each level across animals to facilitate between-group comparison. 2) The final stimulus duration was set at 5 s, reflecting a moderate level of difficulty. This was selected based on pilot data showing that 6-OHDA lesioned animals can reach a performance level comparable to that of control animals at this difficulty level. Differences in reversal learning can thus be interpreted as differences in cognitive flexibility rather than differences in operant learning per se. 3) During reversal phase of training, the rule was switched from rewarding nose poke into a lit aperture to rewarding nose poke into a dark aperture.

**Figure 1.**
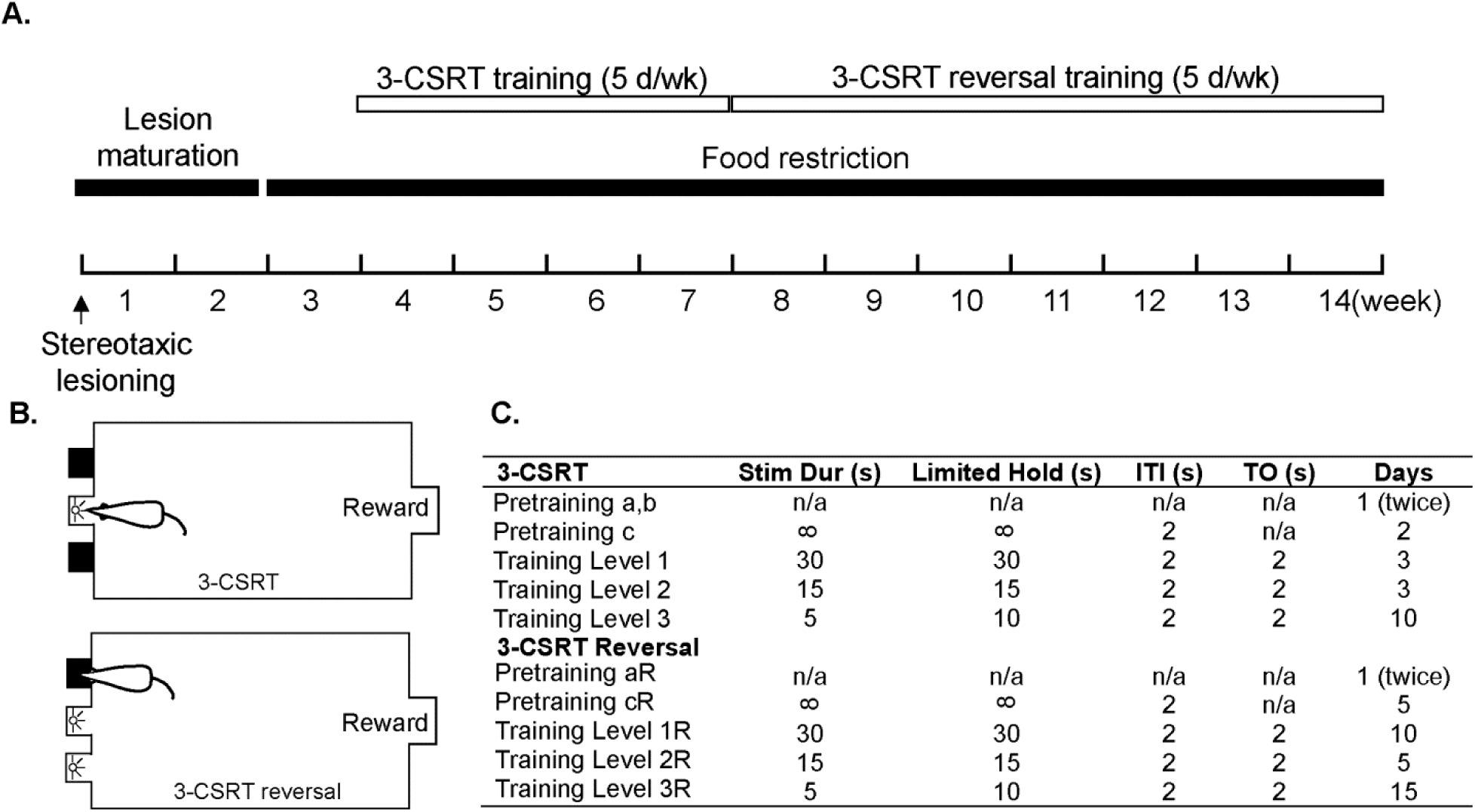
Experimental protocol for operant training. **(A)** Timeline of experiment. **(B)** 3-CSRT and reversal learning. **(C)** Progressive training schedule.

Training was started 1 week following the initiation of food restriction and was always performed between 8 a.m. and 1 p.m. Each operant cage (MedAssociates, St. Albans, VT) consisted of a sound attenuating cubicle (63 cm width, 46 cm depth, 61 cm height) with a fan which was always turned on during testing, modular test chamber (33 cm width, 25 cm depth, 33 cm height) with grid floor, house light, 3-bay nose poke wall, pellet dispenser on the wall opposite the nose poke bays, pellet trough receptacle, receptacle light, head entry detector, smart controller, and infrared camera (Birdhouse Spy Cam, West Linn, OR, USA) for real-time viewing of animals on a TV monitor. Cages were operated by MED-PC software using a personal computer. Pellet dispensers were loaded with dustless sucrose pellets (45 mg/pellet, #F0025, Bio-Serv, Flemington, NJ, USA). Behavioral training was implemented with a fixed ratio 1 schedule response-reward task (up to 90 trials or 30 min/day, 5 days/week). The walls, nose poke apertures, food receptacle, and grid floor were wiped with 70% isopropyl alcohol between animals.

#### Habituation and shaping of behavior

Rats were familiarized with the test chamber and sugar pellets prior to training. Nose poke and reward retrieval behavior were shaped in pretraining. In pretraining ‘*a*’, 10 sugar pellets were put in each nose poke aperture and pellet receptacle, and the animal was allowed to explore the test chamber and retrieve the pellets for 15 min. In pretraining ‘*b*’, the animal was kept in the test chamber for 10 min, while a sugar pellet was dispensed into the receptacle every 20 s with the receptacle light turned on. The receptacle light was turned off 2 s after detection of a head entry (reward retrieval). Pretraining ‘*a*’ and ‘*b*’ were repeated once during the same day. In pretraining ‘*c*’, the animal was trained to associate nose poking into a lit aperture with receiving a single sugar pellet reward. Each daily session lasted 30 min or until the animal received 90 rewards. For each trial, the light in a pseudorandomly chosen nose poke aperture was turned on (stimulus). When a nose poke was detected in the lit aperture (correct nose poke), the aperture light was turned off, and the light in the pellet receptacle was turned on with a sugar pellet dispensed (reward). The receptacle light was turned off 2 s after detection of a head entry. After a 2-s inter-trial interval (ITI), the next trial was started.

#### 3-CSRT Training

During the regular phase of 3-CSRT training, the animal was trained following a progressive schedule (**Fig. 1B, C**). The animal was trained to make a correct nose poke in response to a relatively short stimulus duration. Each single daily session lasted 30 min or until 90 trials were reached. At the start of each trial, the chamber light was turned on and a randomly selected stimulus was started. The stimulus stayed on for a set duration or until a nose poke (correct or incorrect) was detected. The animal received a food reward following a correct nose poke within the set limited hold duration, which was set to be the same as the stimulus duration or slightly longer for short stimulus durations. Following reward retrieval and ITI, the next trial was started. If an incorrect nose poke was detected, the animal was punished with a time out (TO), during which the chamber light was turned off for 2 s. If no nose poke was detected within the limited hold duration, an omission was recorded, and the animal punished with a TO. After each TO, the chamber light was turned on, and after an ITI, the next trial was started. If a nose poke was detected during the ITI, a premature response was recorded without incrementing the trial number, and the animal punished with a TO. Any nose pokes following a correct response and before reward retrieval were recorded as perseverative responses.

#### Reversal training

During the reversal phase of 3-CSRT-R training (**Fig. 1B, C**), the stimulus was switched from a lit aperture among dark apertures to a dark aperture among lit apertures. The animal was trained progressively to learn to nose poke the dark aperture to receive reward.

Analysis of the operant behavior included [7]:

- *nose poke accuracy* = (number of correct responses)/(number of correct + number of incorrect responses) * 100%, a primary measure of operant learning
- *omissions* rate = (number of omissions)/(number of trials completed), a measure of attention
- *premature responses*, a measure of impulsivity
- *perseverative rate = (*number of perseverative responses)/(number of correct responses), a measure of compulsive behavior
- *correct nose poke latency* = average time from onset of stimulus to a correct response, a measure of attention and cognitive processing speed
- *reward retrieval latency =* average time *from correct response to retrieval* of sugar pellet, a measure of *motivation*

### Statistical analysis

Data are presented as the mean ± S.E.M. and analyzed using GraphPad Prism (version 8.3.0, GraphPad Software, San Diego, CA). All data were subjected to the Shapiro-Wilk test for normality. The following data transformations were applied to improve normality: arcsine for nose poke accuracy, logarithm for nose poke latency, reciprocal for reward latency, square root for premature responses and perseverative rate. 3-CSRT data were analyzed using a two-way ANOVA with repeated measures for each training level, with lesion and time as the factors, and with Holm-Sidak’s multiple comparisons test. Data for individual days that failed the normality test were excluded from ANOVA test and analyzed separately using the Mann-Whitney test. Data were subjected to Bartlett’s test for homogeneity of variance for each training level. Data that failed the Bartlett’s test were analyzed using Student’s or Welch’s *t*-test (based on Levene’s test for homogeneity of variance) to compare 6-OHDA lesioned and control group on individual days. Overnight activity data were analyzed using paired Student’s *t*-test. Accelerating rotarod and sucrose preference data were analyzed using unpaired Student’s *t*-test. *p* < 0.05 was considered statistically significant.

## Results

### Lesion verification

TH immunostaining confirmed that the dopamine-depletion lesion was mainly limited to the dorsomedial aspect of the striatum, a region of the basal ganglia central to cognitive processing. **Fig. 2B** shows part of a representative brain slice at bregma + 0.72 mm with reduced TH immunoreactivity in the dorsomedial striatum in the right hemisphere (compared to dorsolateral and ventral striatum, and to control **Fig. 2A**). There was also loss of TH-immunoreactive cells in the substantia nigra pars compacta (**Fig. 2D** compared to **2C**) induced by 6-OHDA lesioning of the dorsomedial striatum. Lesioned striatal area was quantified as a percentage of total striatal area bilaterally at three representative bregma levels designated as rostral, mid and caudal striatum (**Fig. 2E**), which showed (21.32 ± 4.09 %), (32.37 ± 2.39 %), and (34.93 ± 2.59 %) lesion, respectively.

**Figure 2.**
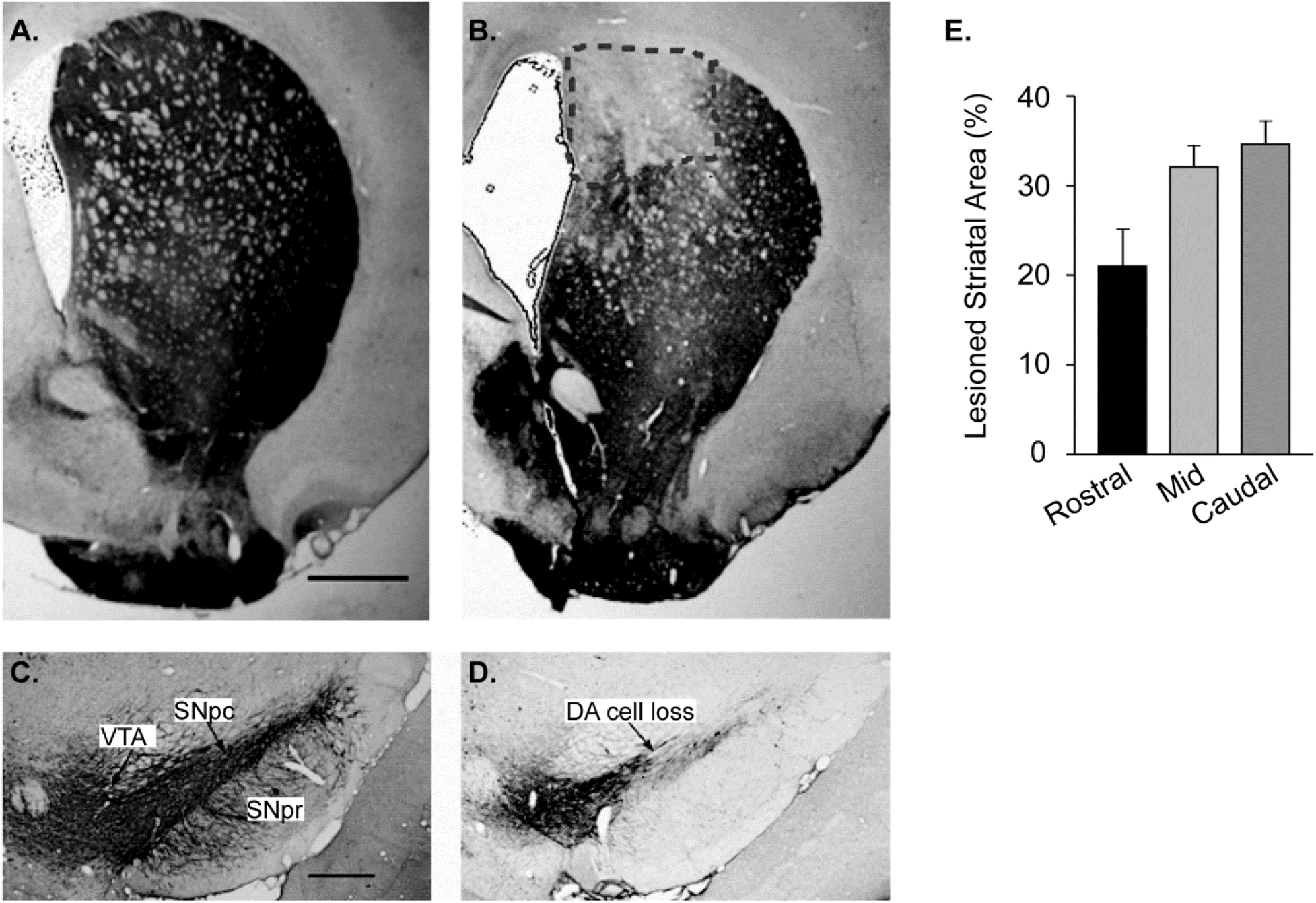
Immunostaining for tyrosine hydroxylase to determine the degree and anatomical site of lesion. Representative images of coronal sections reveal a loss in TH immunoreactivity in the striatum **(B**, at bregma + 0.72 mm) and substantia nigra pars compacta **(D**, at bregma -5.28 mm) in a 6-OHDA lesioned animal compared to a control animal **(A, C). (E)** Lesioned striatal areas are quantified as percent of bilateral striatal area at rostral, mid, and caudal levels (*n* = 8). Scale bar = 1 mm in A, B; 0.5 mm in C,D.

### Lesion spared motor functions and sucrose preference

Overnight activity measurement in home cage showed a trend of decreased horizontal activity counts two weeks after 6-OHDA lesioning (15,077 ± 1,296 counts, *n* = 13) compared to baseline (18,223 ± 1,815 counts, *p =* 0.074, paired Student’s *t*-test), but no difference in vertical activity counts at two weeks (4,870 ± 1,129 counts) compared to baseline (6,033 ± 1,531 counts, *p =* 0.52. **Fig. 3A**). There was also no significant lesioning effect on the maximum velocity during any 15-minute intervals (data not shown). In the accelerating rotarod test, no significant differences were evident in latency to fall between control (169 ± 15 s, *n* = 8) and lesioned animals (161 ± 22 s, *n* = 8, *p =* 0.77, unpaired Student’s *t*-test, **Fig. 3B**). Analysis of sucrose preference (**Fig. 3C**) revealed that 6-OHDA lesioned animals (74.76 ± 4.60 %, *n* = 8) did not differ from controls (78.76 ± 6.06 %, *n* = 6, *p =* 0.20, unpaired Student’s *t*-test).

**Figure 3.**
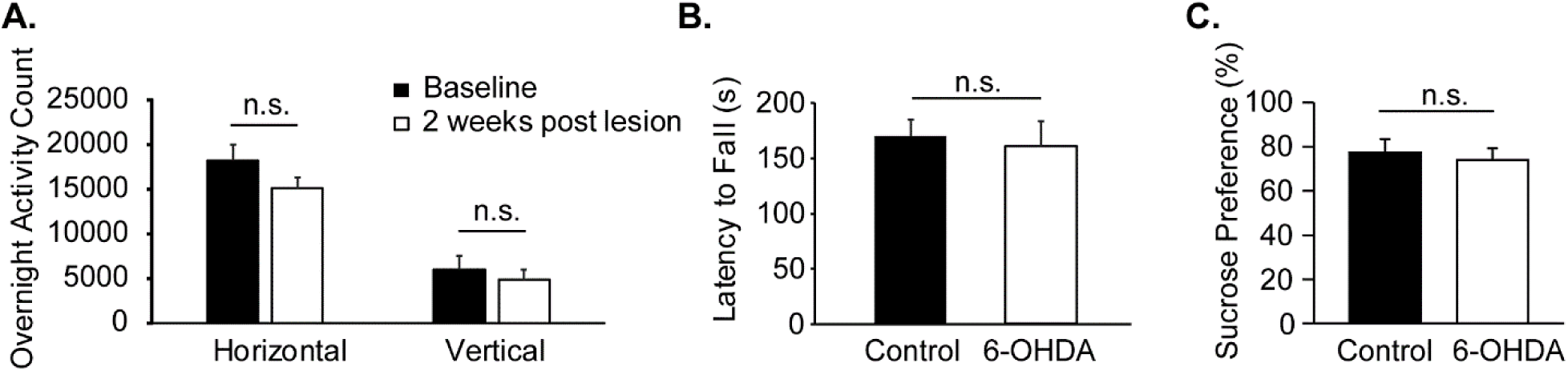
Lesion largely spared motor functions and sucrose preference. **(A)** Overnight locomotor activity before and 2 weeks after 6-OHDA lesioning (*n* = 13). **(B)** Accelerating rotarod test showed no significant differences in mean latency to fall between lesioned and control rats (*n* = 8). **(C)** Analysis of sucrose preference revealed no significant differences between lesioned (*n* = 8) and control rats (*n* = 6, *p =* 0.20, unpaired Student’s *t*-test).

### Lesion induced severe deficits in the reversal learning phase of 3-CSRT

During the acquisition phase of 3-CSRT (Levels L1, L2, L3), both lesioned and control rats showed improvement in nose poke accuracy and shortening of correct nose poke latency (**Fig. 4**). Lesioned rats compared to controls showed: 1) statistically significant lower nose poke accuracy (*p* < 0.001, two-way ANOVA repeated measure) that diminished towards the end of L3 (*p =* 0.16 for day 9 and day 10 of L3, Holm-Sidak test. **Fig. 4A**); 2) a trend of higher omission rate (*p =* 0.061 for day 2 of L1 and day 1 of L2, Mann-Whitney test) with statistically significant differences in variance (Levene’s test. **Fig. 4B**); 3) a trend of higher correct nose poke latency in L1 (*p =* 0.070) and L2 (*p =* 0.088) that diminished in L3 (*p =* 0.19, ANOVA. **Fig. 4C**); 4) statistically significant greater reward retrieval latency (**Fig. 4D**); 5) no differences in premature responses (**Fig. 4E**) and fecal pellet count (**Fig. 4G**); and 6) statistically higher perseverative rate in L1 (*p =* 0.033, ANOVA) and day 1 of L2 (*p =* 0.043, Mann-Whitney) that diminished in L3 (*p =* 0.25, ANOVA. **Fig. 4F**).

**Figure 4.**
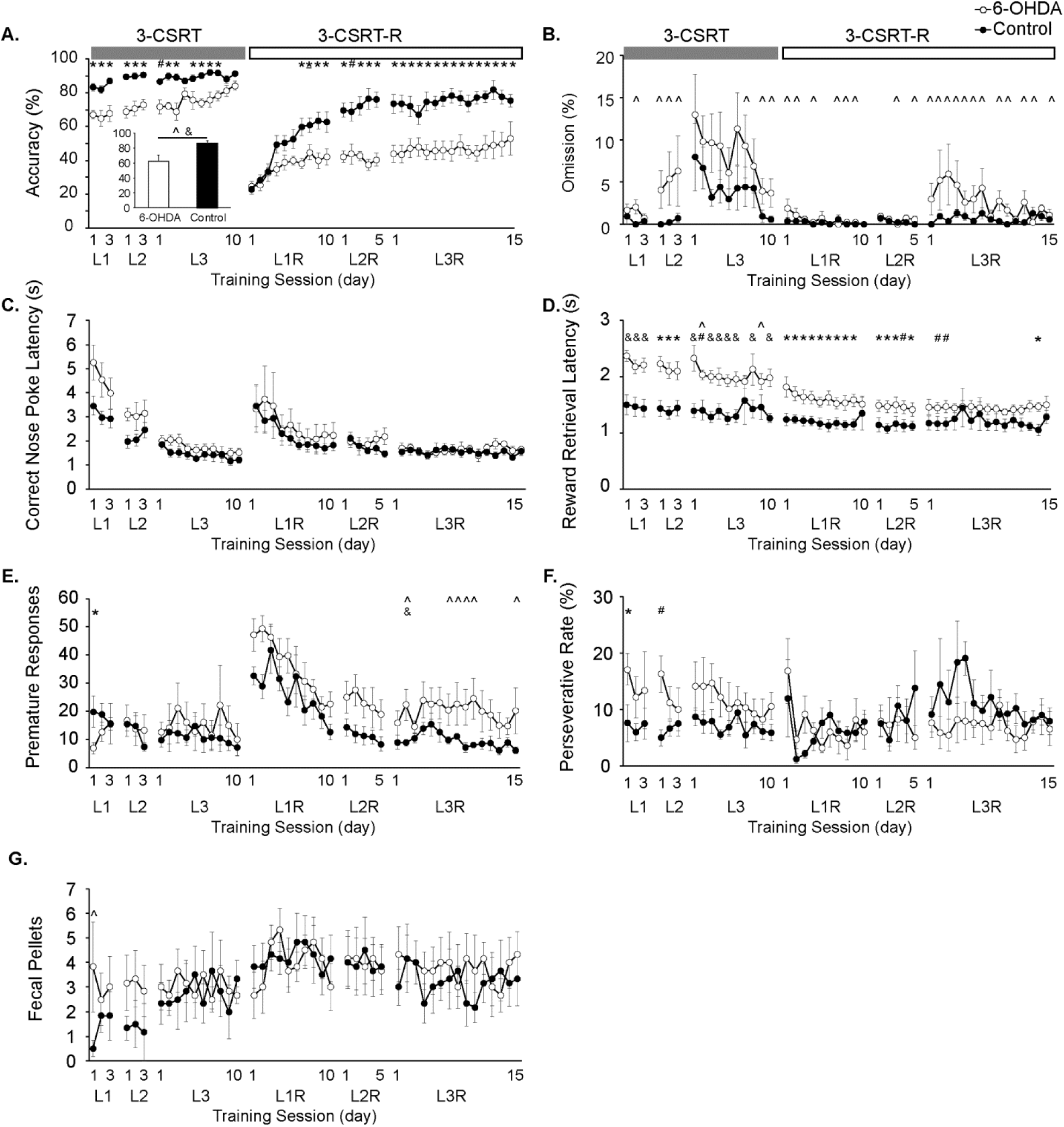
Differences in 3-CSRT acquisition and reversal learning (3-CSRT-R) in 6-OHDA lesioned animals (*n* = 6) compared to controls (*n* = 6). **(A)** While 6-OHDA animals were moderately impaired in 3-CSRT acquisition with lower nose poke accuracy, profound deficits were noted during the reversal phase compared to controls. Inset shows average nose poke accuracy over the last five days of L3R (level 3, reversal) normalized by the mean of last five days of L3 (level 3). The normalize accuracy was significantly lower in 6-OHDA animals (*P* < 0.05, *t*-test). **(B)** Omission rate. **(C)** Correct nose poke latency. **(D)** Reward retrieval latency was longer in 6-OHDA animals compared to controls. **(E)** Premature responses. **(F)** Perseverative rate. **(G)** Fecal pellet count. *P* < 0.05, 6-OHDA vs. control groups: * Holm-Sidak *post hoc* test, # Mann-Whitney test, & Student’s or Wilch’s *t*-test, and ^ Levene’s test for homogeneity of variance.

At the start of 3-CSRT-R, when the rule for correct (rewarded) response was switched from nose poking a lit aperture to nose poking a dark aperture, both lesioned and control rats showed sudden drop in performance with decreased nose poke accuracy to the same extent, increased correct nose poke latency, and increased premature responses. Both groups showed improvement in these parameters with continued training. Control rats improved nose poke accuracy rapidly to a plateau of about 75%, while lesioned animals only improved accuracy modestly to a plateau of about 50%. There were statistically significant differences between the two groups (**Fig. 4A**, L1R, *p =* 0.029; L2R, *p =* 0.0003; L3R, *p =* 0.0025. ANOVA). **Fig. 4A** inset shows average nose poke accuracy over the last five days of L3R normalized by the mean of last five days of L3. The normalize accuracy was significantly lower in 6-OHDA animals (*P* < 0.05, *t*-test). Lesioned compared to control rats continued to demonstrate significantly greater reward retrieval latency (**Fig. 4D**, L1R, *p =* 0.0075; L2R, *p =* 0.027; L3R, *p =* 0.052. ANOVA). A nonsignificant trend of higher premature responses was noted in lesioned compared to control rats during the later stages of reversal training (**Fig. 4E**, L1R, *p* = 0.10; L2R, *p* = 0.058, ANOVA. L3R, day 2, *p =* 0.037, Welch’s t-test) with significant differences in variance (L3R, Levene’s test). Also noted were significant differences in variance in omission rate between the lesioned and control rats (**Fig. 4B**, Levene’s test). No between-group differences were noted in correct nose poke latency, perseverative rate, and fecal pellet count.

## Discussion

We modified the well-established 5-CSRT paradigm to 3-CSRT with nose poke and reversal learning to test cognitive flexibility in animals with 6-OHDA lesion to the dorsomedial striatum. Lesioned animals compared to controls showed robust deficit in reversal learning, in the absence of anhedonia and general deficits in motor functions, operant learning, attention, and impulse control.

### Acquisition of 3-CSRT

The CSRT paradigm is an operant learning task widely used to study attention and impulse control in rodents [6]. The task requires a consecutive series of information processing, decision making, and actions, including: waiting for the stimulus and inhibition of premature responses during the inter-trial interval → attention to the stimulus → recall of prior successful responses → choice of nose poke response → nose poke → recall of reward retrieval → decision to initiate reward retrieval → reward retrieval. Control rats quickly learned the 3-CSRT task and reached a plateau of about 90% accuracy. Lesioned rats, while showing deficits in the initial phase of training, were able to reach a comparable level of accuracy (84.24 ± 2.64%) after 16 days of training. During the 16-day acquisition of the 3-CSRT, there were no significant group differences in premature responses, except on day 1 when control animals demonstrated greater premature responses. This suggests that during acquisition there was little evidence for a group difference in impulsive behavior. The lesioned animals compared to controls showed a trend of higher omission rate as well as statistically significant differences in standard deviation, suggesting a mild lesion-induced deficit in attention.

Lesioned compared to control rats demonstrated a significantly greater reaction time for reward retrieval. Although motor deficits could in principle contribute to group differences seen in reward retrieval latency and omission rate, several lines of evidence argued against general motor impairment. There were no significant group differences in spontaneous locomotor activity or in general motor strength, balance and coordination as measured using the accelerating rotarod test. Of importance, the correct nose poke latency was almost identical between the two groups, suggesting comparable level of attention, speed for information processing and decision making, and speed to nose poke action. Likewise, no lesion effect was noted in the appetitive preference for sucrose reward using the sucrose preference test. This suggests that differences in reward retrieval latency (or omissions rate) likely reflect a slowing of cognitive processing of reward expectation and mildly impaired attention, rather than general motor dysfunction, lack of motivation, or severe attention deficit. The number of fecal pellets counted during the learning phase showed no group differences, suggesting no lesion effect on anxiety-like behavior during the cognitive challenges in 3-CSRT.

### Reversal learning in 3-CSRT

The reversal learning phase was initiated at a time point when lesioned and control animals reached similar level of accuracy. During the initial stage of reversal learning (3-CSRT-R), both lesioned and control rats showed a sudden drop in nose poke accuracy to the same extent. However, with continued training of only a few sessions, control rats rapidly improved their performance, reaching a plateau of about 80% accuracy, while lesioned animals only improved modestly and reached a plateau of about 50% accuracy. We further normalized the average accuracy over the last 5 days of reversal learning (L3R) by the mean of accuracy over the last 5 days of regular training (L3), to control any possible lesion-related deficits in motor and cognitive functions. The normalized 3-CSRT-R accuracy was significant lower in lesioned (61.83 ± 8.78 %) compared to control animals (86.00 ± 4.28 %) (**Fig. 4A** inset). Thus, the 3-CSRT-R task unmasked lesion-induced deficits in cognitive flexibility.

Deficits in reversal learning can be impacted by ‘perseverant’ responses, that is, the inappropriate maintenance of responses previously associated with either reward (learned-reward response) and/or with non-reward (learned-nonreward responses) [14]. The exact contribution of persistent learned-reward or learned-nonreward responses in the lesioned animals, and the role these might play in inhibiting new learning of the reversal task is unclear. There was trend of greater premature responses and omission rate in lesioned compared to control animals, suggesting mild impairment in impulse control and attention.

Our findings suggest that learning is substantially more rapid in the 3-CSRT and 3-CSRT-R nose-poke tasks than has been typically reported with either the 5-CSRT and 5-CSRT-R lever-press task [15,16], the 5-CSRT touchscreen task [17], or the 5-CSRT nose-poke task [6,7]. In part, such difference may be related to the fact that nose-poke responses occur at a higher baseline rate compared to those of lever pressing or touchscreen responses. Significant lesion effect in nose poke accuracy was achieved with a relatively small number of animals (*n* = 6/group), possibly a reflection of smaller variability in a behavior well within the natural repertoire of the animal.

Of note, De Bruin et al. previously applied a variant of the 5-CSRT lever as a 2-CSRT lever pressing task [18], a paradigm later adapted by Homberg et al. to a two-choice nose-poke paradigm (2-CSRT) [19]. The latter, using a fixed-ratio 3 schedule of reinforcement in nonlesioned rats, demonstrated learning acquisition in 25 training sessions of 50 trials per session, and reversal learning to criterion performance in 3 sessions. This shortened duration for reversal training is consistent with the notion that reversal learning decreases in difficulty as the number of holes available for nose poke decreases. While the shortening of training time is desirable, the lower level of task difficulty may decrease the sensitivity to detect deficits in executive function. Therefore, experimental design, and the choice of 5-CSRT or 3-CSRT should be based upon the anticipated magnitude of deficit.

### Dorsomedial striatal lesions

An early feature of PD is deficits in cognitive flexibility, an aspect of executive functions, which involve cognitive processes of set-shifting, working memory, and information processing. Dopamine loss in PD patients is predominant in the posterior putamen, a region associated with the control of habitual behavior. It has been proposed that executive dysfunction, may result as patients become overly reliant on the goal-directed mode of action control that is mediated by comparatively preserved processing in the rostromedial striatum [20]. While not identical to PD, the 6-hydroxydopamine lesioning of dopaminergic neurons of the nigrostriatal system reproduces many of its features [10]. Dopaminergic lesions in the dorsomedial striatum of rodents are critical for successfully observing impaired reversal learning [3,21,22]. Past work has shown that the formation of the critical action-outcome associations mediating goal-directed learning are localized to the dorsomedial striatum, whereas the sensorimotor connections that control the performance of habitual actions or procedural learning are localized to the dorsolateral striatum [23]. In patients and in the animal models, such deficits presumably reflect alterations in frontostriatal processing [12]. Our TH immunostaining results showed that lesions were primarily localized in the mid-caudal levels of the striatum, and appropriately limited to the dorsomedial quadrant of the striatum. Retrograde dopaminergic cell losses were also apparent bilaterally in the substantia nigra. While in 6-OHDA rodent model deficits in cognitive flexibility have previously been examined using the cross-maze [3], T-maze [24], and a food-digging task [25], the current study is the first to examine such deficits during the reversal phase of a choice serial reaction time task. In fact, the most dramatic and persistent learning deficits in lesioned animals were noticed during the reversal learning phase. Our results demonstrate that deficits in cognitive flexibility can be robustly detected in the 3-CSRT-R nose-poke paradigm after two weeks of reversal training. While differences in experimental design and criteria for determining ‘learning’, as well as rodent strains may affect the final duration of experimentation, our findings underscore the practicality of using a 3-CSRT-R nose poke paradigm in evaluating cognitive flexibility. We propose that the 3-CSRT-R testing in rats with bilateral dorsomedial striatal lesions may be a useful model for future testing of treatments aimed at improving executive dysfunction in PD.

## Acknowledgments

This work was supported by grants from the US Department of Defense (Army, CDMRP) grant # W81XWH18-1-0666 (DPH) and grant # W81XWH19-1-0443 (MWJ). We thank Dr. Giselle M. Petzinger for constructive discussion about the project.

